# Regimen-dependent glucocorticoid effects improve muscle performance without altering CNS physiology in *mdx* mice

**DOI:** 10.64898/2026.03.11.711227

**Authors:** Gretel S. Major, Jiayi Chen, Eva van den Berg, Deirdre Merry, Angus Lindsay

**Affiliations:** School of Biological Sciences, University of Canterbury, Christchurch 8041, New Zealand; Biomolecular Interaction Centre, School of Biological Sciences, University of Canterbury, Christchurch 8140, New Zealand; Department of Medicine, University of Otago, Christchurch 8014, New Zealand; Maurice Wilkins Centre for Molecular Biodiscovery, Auckland, New Zealand

## Abstract

Duchenne muscular dystrophy (DMD) is a multisystem disorder affecting striated muscle, metabolism, and the central nervous system (CNS). Although glucocorticoids remain the standard therapy, muscle-centric evaluations typically fail to capture how dosing regimen and compound selection affect CNS and metabolic phenotypes. Here, we compared daily and weekly dosing of prednisolone and vamorolone in juvenile *mdx* mice over six weeks to determine how these variables influence multisystem outcomes. Multiorgan efficacy and adverse effects were quantified across behavioural, endocrine, metabolic, cardiovascular, and muscle domains using behavioural assays, *in vivo* and functional muscle testing, haemodynamic evaluation and histopathology. Daily glucocorticoid dosing failed to improve muscle function or strength, whereas weekly vamorolone produced the most robust improvements in functional and *in vivo* muscle strength. Daily prednisolone reduced circulating creatine kinase levels, but this biochemical change did not translate into enhanced muscle function outcomes. Daily regimens also induced severe adrenal cortical atrophy, yet these endocrine alterations were dissociated from CNS stress and anxiety responses, which remained unchanged by treatment. In addition, daily dosing caused pronounced systemic metabolic consequences, whereas weekly regimens substantially attenuated these effects, identifying dosing frequency as a key determinant of safety. Together, these findings demonstrate that glucocorticoid regimen selection fundamentally reshapes the efficacy–adverse effect profile and underscores the value of integrated multiorgan evaluation in DMD. This work highlights the need to expand therapeutic assessments beyond muscle pathology and raises new questions about how glucocorticoid signalling differentially engages peripheral and central physiological systems.

## 1 Introduction

Duchenne muscular dystrophy (DMD) while conventionally thought of as purely a severe muscle wasting condition, is becoming more commonly understood as a life-limiting, multi-system physiological disorder with central nervous system (CNS) involvement (Vaillend *et al*., 2025). Pathogenic variants within the *DMD* gene locus, which encodes the cytoskeletal protein dystrophin, result in reduced expression of multiple dystrophin isoforms across striated muscle and the CNS (Hoffman *et al*., 1987). Loss of dystrophin disrupts fundamental physiological processes including force transmission (Hughes *et al*., 2017), membrane stability (Goldstein & McNally, 2010), and excitation-contraction coupling in skeletal and cardiac muscle (Ullrich *et al*., 2009), culminating in progressive muscle weakness and cardiorespiratory insufficiency. In parallel, dystrophin deficiency within the CNS alters neural circuit function (Vaillend & Chaussenot, 2017; Caudal *et al*., 2020), stress responsivity (Razzoli *et al*., 2020; Maresh *et al*., 2023), and behavioural regulation (Saoudi *et al*., 2021), giving rise to complex neurocognitive and neurobehavioural comorbidities (*e.g.,* ADHD, autism spectrum disorder features, lower IQ) which coexist with primary muscle pathology (Darmahkasih *et al*., 2020). Collectively, these manifestations reflect a system-wide disturbance of physiological homeostasis rather than isolated tissue pathology, necessitating evaluation of therapeutic efficiency across multiple physiological domains, including skeletal muscle, the CNS and the cardiovascular system.

Glucocorticoids (GCs), including prednisolone, remain the only standard-of-care therapy for treating DMD, slowing disease progression and prolonging ambulation (Bello *et al*., 2015). GCs are proposed to delay DMD progression predominantly through anti-inflammatory mechanisms that attenuate muscle degeneration (Angelini & Peterle, 2012). However, GC receptor activation exerts broad physiological effects across multiple organ systems, so chronic exposure is fraught with complexity and systemic burden, including impaired growth, osteoporosis, obesity, metabolic dysregulation, behavioural changes and suppression of the hypothalamic-pituitary-adrenal (HPA) axis (Handberg *et al*., 2022). These adverse effects reflect dose- and regimen-dependent disruption of endocrine, musculoskeletal, cardiovascular and neurobehavioural physiology which substantially compromises patient quality of life. Clinical evidence suggests that intermittent GC dosing may preserve or enhance muscle function relative to daily administration (Connolly *et al*., 2002; Bello *et al*., 2015), with supporting preclinical studies at lower GC doses (Quattrocelli *et al*., 2017). Moreover, newer dissociative steroids such as vamorolone demonstrate reduced systemic side effects while retaining anti-inflammatory efficacy (Heier *et al*., 2013), yet their impact on integrated physiological function, particularly cardiac haemodynamics and CNS-related outcomes, remains incompletely characterised, and they have not been evaluated under intermittent dosing regimens.

Given that neurological manifestations occur in approximately half of individuals with DMD (Darmahkasih *et al*., 2020) and are associated with poorer clinical outcomes (Mochizuki *et al*., 2008), it is critical to understand how GC compounds and dosing regimens affect CNS functional outcomes within an integrated framework that also considers muscle structure and function, cardiovascular physiology and systemic metabolism. This six-week study directly compared daily and weekly prednisolone (5 mg/kg/day) and vamorolone (30 mg/kg/day) in the juvenile *mdx* mouse model of DMD during rapid growth and active pathology, using micropipette-guided dosing to minimise stress-axis activation and preserve physiological accuracy (Scarborough *et al*., 2020; Ferreira-Duarte *et al*., 2024). Therapeutic efficacy was assessed across skeletal muscle, cardiac, and CNS phenotypes, alongside stress-axis and endocrine side effects. By integrating multi-organ physiological endpoints with multivariate analyses, this study defined how GC regimen and compound selection shape whole-body physiological function in dystrophin deficiency, highlighting the importance of incorporating cardiovascular and neurophysiological measures into preclinical therapeutic evaluation for DMD.

## 2 Materials and methods

### 2.1 Ethical approval

Ethical approval for the research protocol and all procedures was obtained from the University of Otago Animal Ethics Committee (AUP-24-122). All procedures were performed in accordance with the Animal Welfare Act and the ARRIVE guidelines for animal experimentation (du Sert *et al*., 2020).

### 2.2 Animal breeding and husbandry

Dystrophin-negative C57BL/10ScSn-*Dmd^mdx^*/J (*mdx*) and dystrophin-positive C57BL/10ScSnJ (WT) mice were bred in-house at the Christchurch Animal Research Area (AUP-23-115) from stock sourced from Jackson laboratories (Bar Harbor, Maine and Sacramento, California, USA) on a 12-hour light/12-hour dark cycle, 20°C–25°C, 40% humidity. Pups were weaned at three weeks of age and randomly assigned into cages of 3-4 mice based on treatment group. Body mass was measured every two days, and food consumption was recorded weekly for the duration of the study.

### 2.3 Treatment protocol

*Mdx* mice were treated orally with GCs in daily or weekly regimens for six weeks from four to 10 weeks of age, capturing the period where muscle degeneration peaks and aligning with treatment windows in previous studies **(Figure 1)** (Ziemba *et al*., 2021; Timpani *et al*., 2023; Kourakis *et al*., 2024). Five treatment groups were evaluated: 1) cherry syrup vehicle, 2) 5 mg/kg prednisolone daily, 3) 5 mg/kg prednisolone weekly, 4) 30 mg/kg vamorolone daily and 5) 30 mg/kg vamorolone weekly. WT mice were used as a healthy control receiving only the cherry syrup vehicle. Animals were weighed at 7-8 am every two days, and individual treatments were prepared daily in cherry syrup vehicle relative to body mass to give a final dosage of either 5 mg/kg/d prednisolone or 30 mg/kg/d vamorolone. Mice receiving GC treatments on once weekly regimens received the cherry syrup vehicle on the remaining six days. The amount of cherry syrup vehicle administered was standardised across all treatment groups to achieve 1 uL/g body mass/d. These dosages are consistent with previous preclinical studies evaluating vamorolone (Heier *et al*., 2013, 2019; McCormack *et al*., 2023; Liu *et al*., 2024) and prednisolone (Liu *et al*., 2024; Kourakis *et al*., 2024).

**Figure 1.**
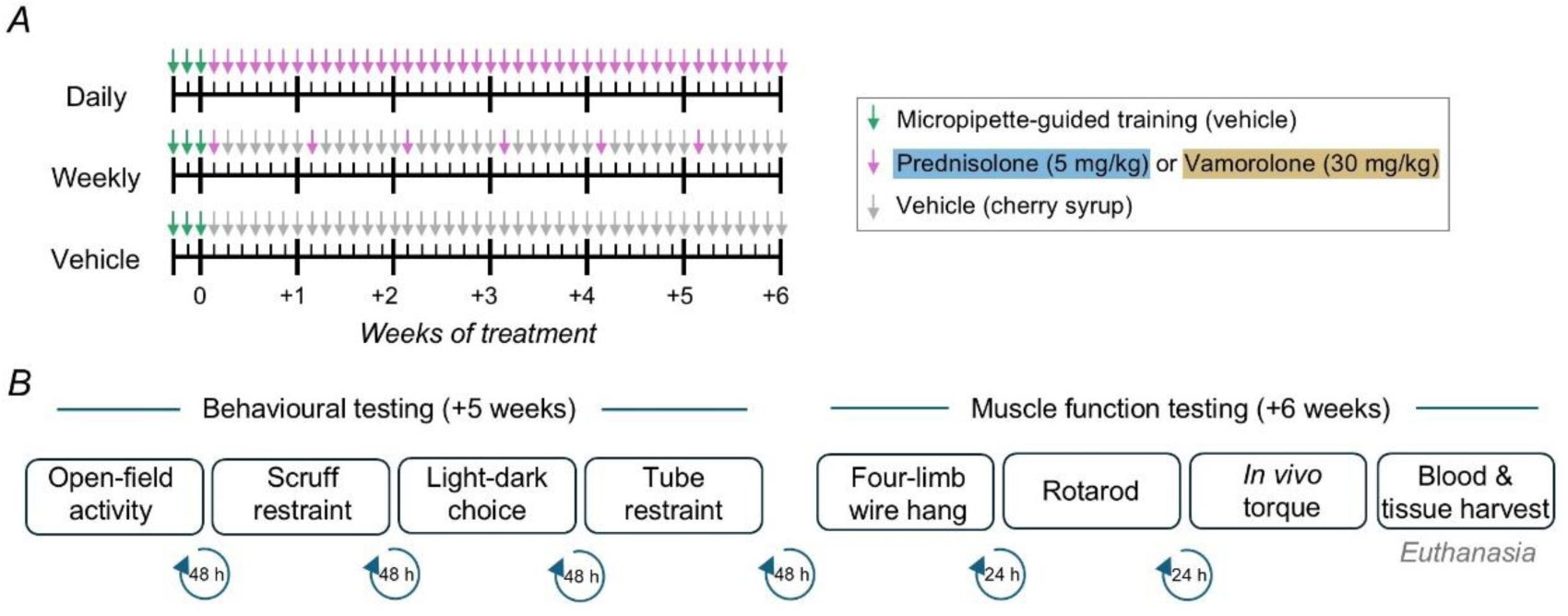
Overview of the treatment regimen and functional assessment timeline. (A) Treatment schematic: Daily or weekly oral dosing of prednisolone (5 mg/kg/day) or vamorolone (30 mg/kg/day) was administered over a 6-week treatment period, alongside vehicle-treated *mdx* and wild-type controls. All treatments were delivered via voluntary micropipette-guided administration following a 3-day acclimation period to minimise stress-axis activation. Weekly groups received the full dose once per week with vehicle on remaining days, whereas daily groups received compound-matched doses each morning. Mice continued on their assigned treatment regimen until euthanasia. (B) Behavioural and muscle-function assessment timeline: Testing batteries were conducted after 5 and 6 weeks of treatment. Behavioural assessments were spaced by 48 h intervals, while muscle-function assessments were spaced by 24 h intervals. Immediately following *in vivo* torque assessments, mice were euthanised and blood and tissues were collected.

Given that *mdx* mice demonstrate stress hypersensitivity in response to a scruff restraint (Lindsay & Russell, 2023; Gharibi *et al*., 2024) and that a primary aim of this study was to determine the effects of GC treatment regimens on behavioural phenotypes in DMD, all treatments were delivered by voluntary micropipette-guided administration. This administration route has been shown to produce similar pharmacokinetic profiles to oral gavage with reduced cortisol production (Scarborough *et al*., 2020; Ferreira-Duarte *et al*., 2024). All mice were trained to drink the cherry syrup vehicle from a 200 μL pipette for three days prior to the beginning of the treatment regimen by positioning a pipette tip containing the cherry syrup vehicle close to the mouth until the mouse drank. All mice were successfully trained after two days and there were no instances of failed drug delivery throughout the study.

### 2.4 Blood glucose

To determine the effects of the treatment regimens on systemic glucose regulation, blood glucose concentrations were measured every two weeks. At week four, mice were fasted for five hours (from ∼8am to ∼1pm) and then blood glucose measured from a tail nick using a blood glucometer (OneTouch Select Plus). For endpoint assessments (after six weeks of treatment), measurements were taken in an unfasted state directly prior to termination.

### 2.5 Behavioural testing battery

The behavioural testing battery was employed after five weeks of treatment (**Figure 1**). To minimise cumulative burden and interaction between behavioural tests, the three tests were conducted 48 h apart. The order of testing was as follows: open-field activity, scruff restraint stress test, light-dark choice, and then tube-restraint stress test with haemodynamic evaluation.

#### 2.5.1 Open-field activity

Total distance travelled in the open-field was measured over 10 min in a plexiglass testing chamber (40 cm L x 40 cm W x 40 cm H, Omnitech) in a room with homogeneous dim illumination (∼50 Lx) using Fusion software (AccuScan, Omnitech Electronics, USA).

#### 2.5.2 Scruff restraint stress test

To evaluate stress hypersensitivity, physical activity was measured for 5 min following a scruff restraint stressor, as described previously (Sekiguchi *et al*., 2009; Razzoli *et al*., 2020; Lindsay *et al*., 2021b) Mice were grasped by the nape between the thumb and index finger, securing the tail between the fourth and fifth finger, and placed in the supine position for 30 s. Following the scruff restraint the mice were immediately placed in a sterile open-field activity monitoring chamber (∼20 cm L x 20 cm W x 40 cm H; Omnitech Electronics, USA). Physical activity was quantified as total movement time over 5 min and reported as both total time and percentage freezing in 1-minute bins (AccuScan, Omnitech Electronics, USA).

#### 2.5.3 Light-dark choice

Anxiety was measured in a light-dark choice apparatus with monitoring over 5 min. The testing apparatus consisted of a plexiglass box with a brightly lit compartment (40 cm L x 20 cm W x 30 cm H; illumination: 600 Lx) connected by an animal entry opening with a trapdoor (10 x 4 cm) to a dark compartment (40 cm L x 15 cm W x 20 cm H; illumination: <10 Lx) (Omnitech Electronics, USA). Each mouse was placed in the dark compartment for 10 s before the trapdoor was opened and mice were allowed to freely explore the apparatus for 5 min. Number of entries and total time spent in the lit compartment were recorded using Fusion software (AccuScan, Omnitech Electronics, USA).

#### 2.5.4 Tube-restraint stress test with haemodynamic measures

The haemodynamic response to stress was measured using a 5 min tube-restraint stressor with concurrent evaluation of haemodynamic parameters. Mice were placed in the CODA^®^ non-invasive blood pressure system using a plexiglass tube mouse holder suitable for the mouse body mass (Kent Scientific, Torrington, CT, USA) for 5 min, as previously described (Gharibi *et al*., 2025). The mouse holders were placed on a 37°C warming platform, and tail cuffs positioned to expose the tip of the tail. Blood pressure was recorded every 30 s for a total of 10 measurements using volume pressure recording sensor technology and CODA software. Mean arterial pressure, heart rate, tail blood volume and shock index (the ratio of maximum heart rate to lowest systolic blood pressure, indicating hypovolemic stress) were recorded. At the completion of the haemodynamic measures, mice were placed into a sterile physical activity monitoring chamber for 5 min to assess physical activity post stressor (as described above).

### 2.6 Muscle function testing battery

Functional muscle strength was tested after six weeks of treatment using the following testing battery: four limb wire hang, rotarod performance and *in vivo* strength of the anterior and posterior crural muscles (24 h between each test; **Figure 1**).

#### 2.6.1 Four-limb wire hang

The four-limb wire hang test assessed whole body strength by measuring the ability of mice to maintain grip on an inverted wire grid mesh (∼1 cm x ∼1 cm mesh) (Treat-NMD protocol DMD_M.2.1.004). The hanging time from mesh inversion was recorded and the test was repeated over three trials for each mouse (15 min between trials). The holding impulse was calculated as body mass multiplied by absolute hang time for the longest trial. One mouse was excluded as they refused the test (hanging < 10 s on 3 repeated attempts).

#### 2.6.2 Rotarod

A rotarod performance test assessed neuromotor coordination over three trials. One day before testing, mice underwent two acclimatisation sessions, 30 min apart, during which they ran for 1 min at 5 rpm before the speed was ramped up to 45 rpm. For the test, each trial started with a stabilisation period at 5 rpm, followed by an acceleration to 45 rpm over 30 s and maintenance at 45 rpm until the mouse fell from the rotarod (or 600 s was reached). Time to fall was recorded for each trial, with the longest trial used for data analysis.

#### 2.6.3 In vivo muscle strength

Maximum isometric strength of the anterior and posterior crural muscles was assessed by stimulation of the common peroneal nerve and sciatic nerve, respectively, using percutaneously placed electrodes connected to a stimulator-force transducer apparatus (Aurora Scientific, Canada). Mice were maintained under anaesthesia (1.5-3% isoflurane, 95% oxygen) throughout and peak isometric tetanic torque was measured by manipulating voltage at 200 Hz every min until a plateau was attained (within 0.01 mN.m). Maximal rate of contraction and maximal rate of relaxation were assessed using peak torque. Torque was analysed as specific torque, *i.e.,* maximum tetanic torque normalised by the combined tibialis anterior (TA) and extensor digitorum longus (EDL) mass for the anterior crural muscles and the gastrocnemius and soleus of the posterior crural muscles.

### 2.7 Creatine kinase

Muscle damage markers were assessed by determination of plasma creatine kinase (CK) levels. A terminal blood sample was obtained following the *in vivo* muscle function tests while the mice were anaesthetised (1.5-4% isoflurane, 95% oxygen). Blood was collected via terminal cardiac puncture into lithium heparin microtubes. Plasma was derived by centrifugation (3,000 g, 5 min, 4°C) and was stored at -80°C until assayed. Creatine kinase levels were quantified spectrophotometrically using a CK-NAC kit as per the manufacturer’s instructions (CK8313, Randox Laboratories).

### 2.8 Tissue harvest

After end-point *in vivo* muscle function testing and blood collection, the organs (brain, adrenal glands, spleen) and striated muscles (diaphragm, TA, EDL, soleus, gastrocnemius and quadriceps) were excised, weighed and snap-frozen in liquid nitrogen for further assays (unless stated otherwise below).

### 2.9 Histopathology and immunohistochemistry

The right TA (opposite leg used for *in vivo* strength testing) and right hemisphere of the diaphragm were coated in OCT (TissueTek) and snap-frozen in liquid nitrogen-cooled isopentane. Muscle samples were thawed to -20°C and cryosectioned at 10 µm. To visualise tissue structure, diaphragm sections were fixed in methanol and stained with haematoxylin and eosin. For quantification of myofibres, mid-belly sections of the TA were fixed in −20 °C acetone for 5 min, blocked in 5% BSA/PBS, and counterstained with laminin (1:500; Sigma-Aldrich L9393) for 2 h at room temperature as previously described (Devananthan *et al*., 2026). Sections were incubated with anti-Rabbit Alexa Flour 488 (1:500; ThermoFisher Scientific A11008) for 1 h at room temperature and mounted in ProLong Golf Antifade with DAPI (ThermoFisher Scientific). Images were acquired on a Zeiss AxioImager Z1 fluorescence microscope and stitched together with Zen software (Zeiss). SMASH software was used to quantify number myofibres, myofibre cross-sectional area and centronucleated fibres (Smith & Barton, 2014). Whole muscle cross sectional area was measured using ImageJ.

The right adrenal gland was immediately fixed in 4% paraformaldehyde for 30 min at RT, coated in OCT and snap-frozen in liquid nitrogen cooled isopentane. Adrenal gland samples were thawed to -20°C and cryosectioned at 4 µm, before staining with haematoxylin and eosin. Images were acquired on a Zeiss AxioImager Z1 microscope and cross-sectional area, cortex area and medulla area measured manually using ImageJ.

### 2.10 Hydroxyproline assay

Muscle fibrosis was assessed by measuring hydroxyproline content of the left hemisphere of the diaphragm according to the manufacturer’s instructions (AB222941, Abcam).

### 2.12 Statistics

All data are reported as mean ± SEM unless otherwise stated. Data were analysed using R and visualised using GraphPad Prism 9. A permutational multivariate analysis of variance (PERMANOVA) was conducted using the Euclidean method to assess overall group differences in multivariate composition across muscle parameters (time to fall during the rotarod test, holding impulse during the four-limb wire hang, maximum isometric torque, creatine kinase levels, centronucleated fibres), behavioural parameters (total distance travelled in the open-field, total movement time following a scruff restraint, total movement time following a tube restraint, time in the lit compartment of light dark choice apparatus, brain mass) or cardiac parameters (blood pressure, heart rate, blood flow, heart mass). One mouse from the *mdx* vehicle control group was removed from the CNS analysis retrospectively, when hydrocephalus was identified. Accordingly, data across all individual tests were standardised prior to PERMANOVA analysis and missing values were imputed (excluding the whole mouse yielded a similar result). For analysis of individual assays, groups of data were compared by one-way ANOVA or Kruskal-Wallis test (when normality assumption was not met), with a P value of <0.05 considered statistically significant. Repeated measures analysis was used for body mass, food consumption and blood glucose analyses. Where relevant, post hoc Tukey’s HSD tests were used for multiple comparisons following ANOVA.

## 3 Results

### 3.1 Prednisolone and vamorolone generate distinct multivariate profiles of muscle functional, contractile and structural physiology

To capture treatment-dependent effects across muscle contractile, structural, and functional parameters, we performed non-metric multidimensional scaling (NMDS) on a combined dataset integrating a subset of physiological outcome measures (four-limb wire hang, rotarod performance, *in vivo* torque testing, plasma CK levels, muscle hydroxyproline content, and the proportion of centrally nucleated fibres; **Figure 2A-B)**. These variables were chosen from the broader dystrophic analysis panel because they reflect complementary aspects of muscle pathology and did not correlate with one another, ensuring that each provided independent physiological information within the integrated analysis. Both GCs, independent of dosing regimen, altered global muscle physiology relative to *mdx* vehicle controls (P ≤ 0.031), with no distinction between daily and weekly prednisolone (P = 0.29), or between daily prednisolone and daily vamorolone (**Figure 2A,B**; P = 0.17). Daily administration of prednisolone produced the largest physiological shift (P = 0.006), followed closely by weekly vamorolone (**Figure 2A,B**; P = 0.009).

**Figure 2.**
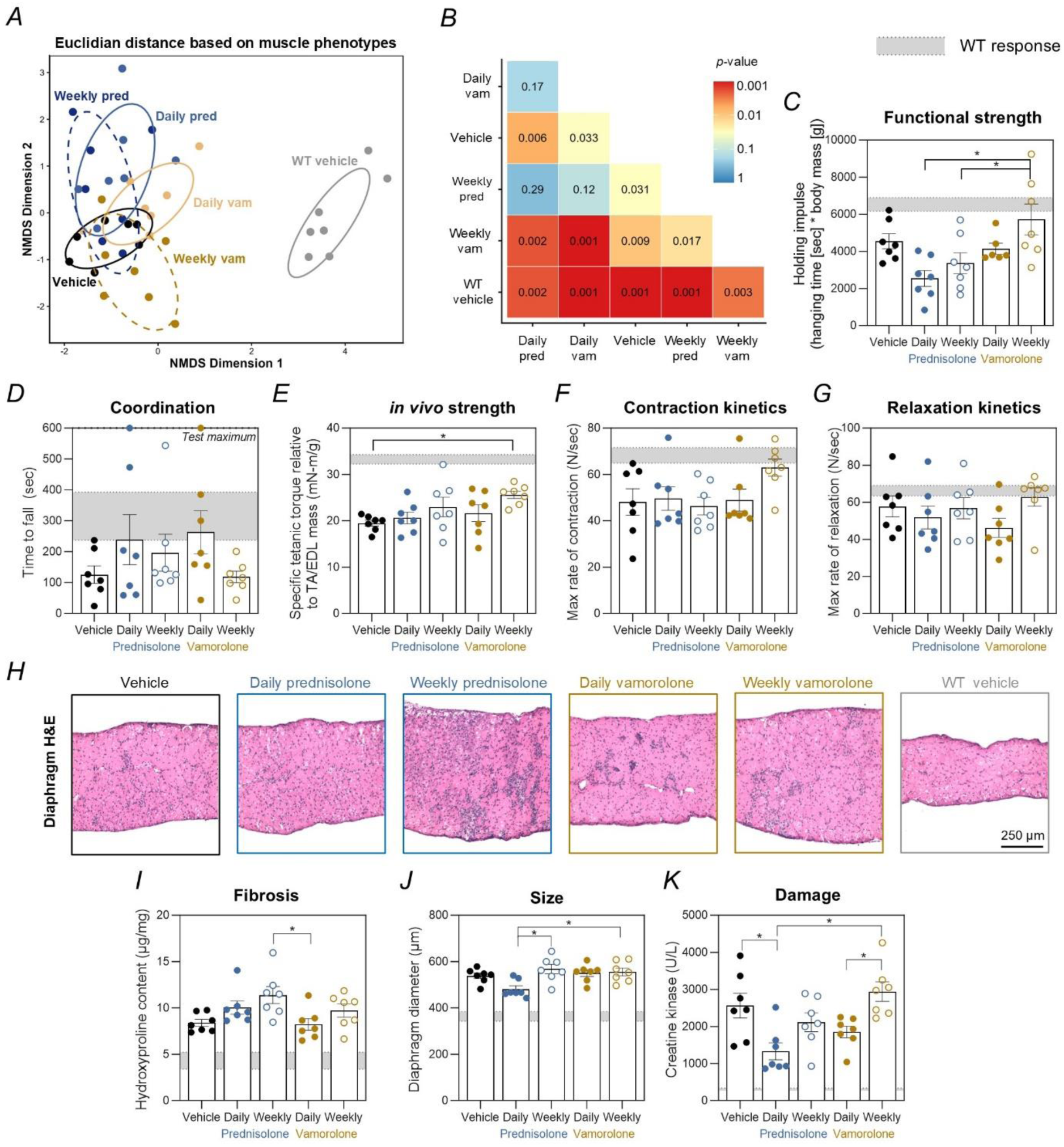
Prednisolone and vamorolone produce divergent profiles of muscle function, contractility, and tissue integrity, reflecting drug-specific physiological effects. (A) Non-metric multidimensional scaling (NMDS) ordination based on Euclidean distance illustrating multivariate skeletal muscle physiological profiles across experimental groups, including prednisolone (blue) and vamorolone (yellow) treatments, and WT (black) and *mdx* (grey) vehicle controls. Each point represents an individual animal, and proximity reflects similarity in integrated physiological profiles. (B) Heat map of pairwise *p*-values for between-group multivariate comparisons across all experimental groups. (C) Four-limb wire hang test assessing functional strength, expressed as holding impulse. (D) Rotarod performance assessing motor coordination, expressed as time to fall. (E) *In vivo* muscle strength expressed as specific tetanic torque normalised to tibialis anterior (TA) and extensor digitorum longus (EDL) muscle mass. (F) Contraction kinetics of the anterior crural muscles expressed as maximal rate of torque development. (G) Relaxation kinetics of the anterior crural muscles expressed as maximal rate of torque relaxation. (H) Representative haematoxylin and eosin (H&E)-stained sections of diaphragm muscle (scale bar = 250 µm). (I) Diaphragm fibrosis quantified by hydroxyproline content. (J) Diaphragm muscle size assessed by muscle diameter. (K) Plasma creatine kinase concentration as a marker of muscle damage. Multivariate group differences were assessed using Euclidean distance–based PERMANOVA on z-score standardised data, with dispersion tested by betadisper. Significant global effects were followed by pairwise multivariate comparisons, and outcome-specific differences were assessed using univariate ANOVA with appropriate correction for multiple testing; *P* < 0.05. WT responses are shown in grey shading as ± SEM. All other data are presented as mean ± SEM; *n* = 7 mice per group.

When evaluating individual physiological outcomes, weekly vamorolone improved functional strength, measured by four-limb holding impulse, relative to both daily and weekly prednisolone (P ≤ 0.029), and was the only group not different from WT vehicle controls (**Figure 2C**; P = 0.87). Rotarod coordination was unaffected by any treatment (P ≥ 0.228, while only weekly vamorolone enhanced *in vivo* strength of the anterior crural muscles relative to vehicle-treated *mdx* mice (P = 0.036), without altering contraction and relaxation kinetics (**Figure 2D-G**; P ≥ 0.20). No treatment improved the strength or contraction rates of the posterior crural muscles (P ≥ 0.094), however, daily vamorolone slowed relaxation relative to prednisolone, weekly vamorolone and vehicle-treated *mdx* controls (**Figure S1A-C**; P ≤ 0.038).

Structurally, no treatment reduced diaphragm fibrosis, as measured by hydroxyproline content, although weekly prednisolone treatment led to greater fibrosis relative to daily vamorolone-treated mice (**Figure 2H,I**; P = 0.042). Daily prednisolone decreased diaphragm thickness relative to weekly dosing of both prednisolone and vamorolone (**Figure 2H,J**; P ≤ 0.036). However, the normalised mass (muscle weight relative to body weight) of the EDL, TA, soleus, or gastrocnemius did not differ among treatment groups relative to *mdx* vehicle controls (**Figure S1D-G;** P ≥ 0.27).

Muscle damage, as indicated by plasma creatine kinase (CK), was reduced only by daily prednisolone relative to vehicle controls (P = 0.008). Weekly vamorolone-treated mice had higher CK than both daily prednisolone- (P = 0.0003) and daily vamorolone-treated mice (P = 0.024), demonstrating that daily GC administration is required to mitigate muscle damage, with prednisolone showing superior efficacy **(Figure 2K)**.

### 3.2 Daily prednisolone alters muscle fibre organisation relative to vamorolone

To further investigate how GC treatment shapes skeletal muscle architecture which underlies contractile performance, we examined muscle fibre organisation. No treatment altered the percentage of centrally nucleated fibres (P ≥ 0.055), indicating comparable levels of ongoing regeneration across groups (**Figure 3A,B**). However, both daily prednisolone and daily vamorolone reduced myofibre cross-sectional area relative to *mdx* vehicle controls (**Figure 3A,C**; P ≤ 0.033). In prednisolone-treated mice, fibre size was also influenced by dosing frequency (P = 0.002), with weekly dosing producing larger fibres than daily dosing (**Figure 3A,C)**. In contrast, vamorolone-treated mice showed consistent fibre sizes across both dosing regimens (**Figure 3A,C**; P = 0.68). Daily prednisolone treatment further increased fibre number per unit area relative to vehicle controls and weekly prednisolone treatment (P ≤ 0.011), an effect not observed with vamorolone treatment (**Figure 3A,D**; P ≥ 0.089). Importantly, structural parameters such as fibre size, density, and central nucleation did not correlate with functional performance measures, rotarod performance, four-limb wire holding impulse, and *in vivo* torque, in *mdx* mice (**Figure S2**), indicating that muscle architecture alone does not predict functional outcomes.

**Figure 3.**
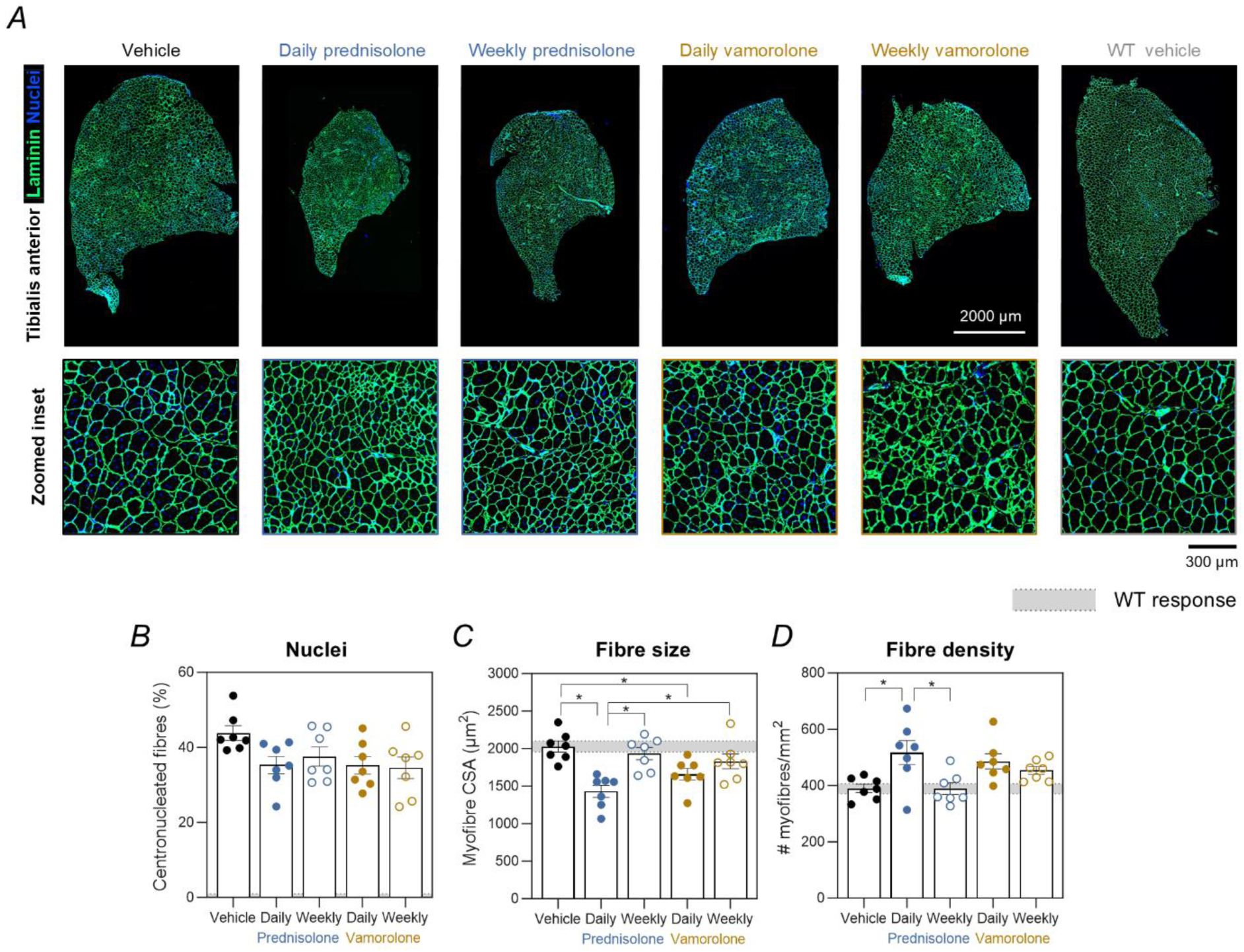
Daily prednisolone enhances fibre number and reduces fibre size, effects less pronounced with vamorolone. (A) Representative laminin (green) and nuclei (blue) immunofluorescence images of tibialis anterior (TA) muscle, showing fibre architecture and myonuclear positioning (scale bar = 250 µm). (B) Percentage of centrally nucleated fibres. (C) Myofibre cross-sectional area (CSA). (D) Fibre density (fibres/mm²). Group differences were assessed using univariate ANOVA; *P* < 0.05. WT responses are shown in grey shading as ± SEM. All other data are presented as mean ± SEM; *n* = 7 mice per group.

### 3.3 Glucocorticoid treatment does not alter brain structure, anxiety or stress-axis physiology

To assess the impact of prednisolone and vamorolone on CNS physiology in DMD, we examined brain size along with behavioural outcomes related to anxiety and stress-reactivity, characteristics of this DMD model. We then performed NMDS to integrate these CNS-related parameters, including responses to scruff-restraint, tube-restraint, light–dark choice, open-field, and brain weight, into a single composite analysis. The NMDS revealed clear separation between vehicle-treated WT and *mdx* mice (P = 0.004), reflecting genotype-dependent differences in overall brain physiology, but no treatment groups diverged from *mdx* vehicle controls (**Figure 4A,B**; P ≥ 0.44). The only distinction observed with *mdx* cohorts was between daily prednisolone and weekly vamorolone groups (**Figure 4A,B**; P = 0.070). Analysis of body mass–adjusted brain mass showed no effect of treatment relative to vehicle controls (P ≥ 0.66), though daily prednisolone resulted in larger brain mass relative to weekly vamorolone (**Figure 4C; Figure S3A,B;** P = 0.040).

**Figure 4.**
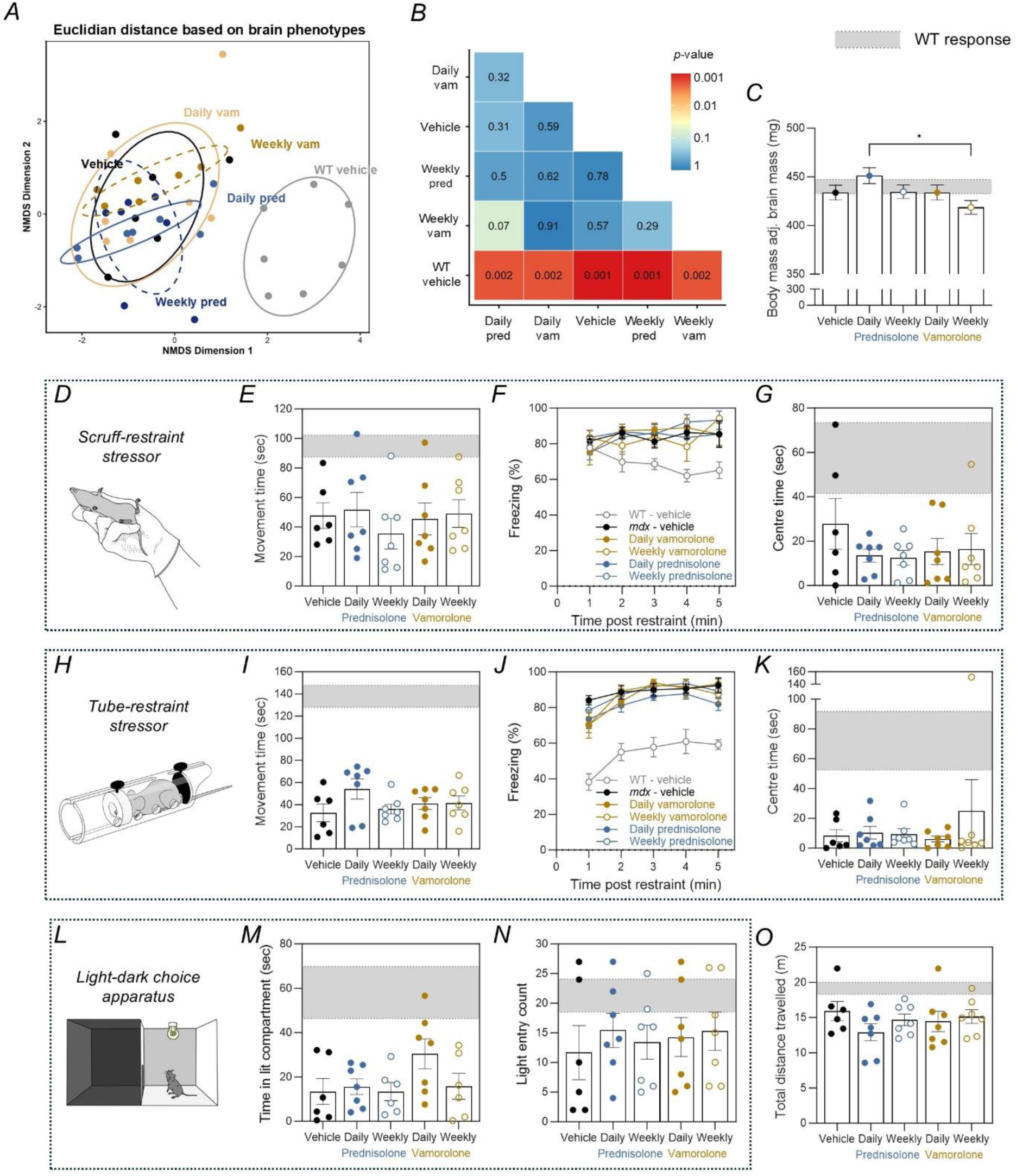
Prednisolone and vamorolone did not alter brain structure or stress- and anxiety-related behavioural outcomes. (A) Non-metric multidimensional scaling (NMDS) ordination based on Euclidean distance illustrating multivariate brain and behavioural profiles across experimental groups, including prednisolone (blue) and vamorolone (yellow) treatments, and WT (black) and *mdx* (grey) vehicle controls. Each point represents an individual animal, and proximity reflects similarity in integrated functional profiles. (B) Heat map of pairwise *p*-values for between-group multivariate comparisons across all experimental groups. (C) Body-weight–adjusted brain mass across treatment groups. Data are estimated marginal means (± SEM) derived from an ANCOVA model with body weight included as a covariate. *Scruff restraint stress:* (D) Schematic of the scruff restraint stress paradigm. (E) Movement time following scruff restraint stress. (F) Percentage of time spent freezing in 1-minute bins. (G) Time spent in the centre of the cage. *Tube restraint stress:* (H) Schematic of the tube restraint stress paradigm. (I) Movement time following tube-restraint stress. (J) Percentage of time freezing in 1-minute bins. (K) Time spent in the centre of the cage. *Light–dark choice anxiety test:* (L) Schematic of the light–dark choice test. (M) Time spent in the lit compartment. (N) Number of entries into the lit compartment. *Open field behaviour:* (O) Total distance travelled over 10 minutes. Multivariate group differences were assessed using Euclidean distance–based PERMANOVA on z-score standardised data, with dispersion tested by betadisper. Significant global effects were followed by pairwise multivariate comparisons, and outcome-specific differences were assessed using univariate ANOVA or Kruskall-Wallis H Test with appropriate correction for multiple testing; *P* < 0.05. WT responses are shown in grey shading as ± SEM. All other data, except where otherwise specified, are presented as mean ± SEM; *n* = 6-7 mice per group.

To probe stress-axis responsiveness, we applied mild (scruff restraint) and moderate (tube restraint) laboratory stressors and tracked movement patterns post stressor **(Figure 4D,H)**. Neither paradigm produced treatment-dependent differences in total movement time (P ≥ 0.23), binned freezing behaviour (P ≥ 0.18), or centre exploration **(Figure 4D-K**; P ≥ 0.95). Importantly, despite previous reports of tonic immobility induced by scruff restraint (Vaillend & Chaussenot, 2017; Lindsay *et al*., 2021a; Lindsay & Russell, 2023), vehicle-treated *mdx* mice were not more sensitive than WT mice (**Figure 4D,E**; P = 0.18). This suggests that repeated daily handling during treatment may have reduced the potency of this stressor, even in the absence of the more intense procedures typically used in the field, such as scruff restraint and oral gavage, possibly reflecting habituation to routine experimental contact.

To evaluate anxiety levels, we used the light-dark choice test and activity in the open field as physiological measures. Treatments did not alter time spent in the lit compartment (P ≥ 0.35), latency to enter (P > 0.99), or number of light entries in the light-dark choice apparatus (**Figure 4L-N**; **Figure S3C**; P > 0.99). Total distance travelled in the open field was also unaffected by treatment (**Figure 4O**; P ≥ 0.45). Collectively, these data indicate that neither prednisolone nor vamorolone significantly modifies brain size or behavioural responses to stress and anxiety stimuli suggesting that short-term GC administration does not exacerbate or alleviate CNS physiological dysfunction in *mdx* mice.

### 3.4 Daily glucocorticoid dosing drive extensive adrenal cortical atrophy

Given the widespread impact of GCs on endocrine systems, and the central role of the HPA axis in stress regulation, we next examined adrenal morphology to assess whether treatment altered peripheral endocrine physiology, independent of behavioural or brain structural changes. Daily dosing of both prednisolone and vamorolone reduced overall adrenal mass (P < 0.0001) and cross-sectional area (P ≤ 0.012) relative to vehicle controls, with daily administration resulting in lighter glands than weekly dosing (**Figure 5A-C**; **Figure S4A,B;** P < 0.0001). This effect was driven by selective atrophy of the adrenal cortex, as absolute medullary size remained unchanged across groups (**Figure 5A,D**; **Figure S4C,D;** P ≥ 0.75). Among all treatments, daily prednisolone caused the most pronounced cortical reduction, resulting in smaller cortical area than any other group (P ≤ 0.028). However, weekly prednisolone (P = 0.032) and daily vamorolone (P < 0.0001) also caused cortical atrophy, whereas weekly vamorolone preserved cortical size relative to vehicle controls (**Figure 5A,E**; P = 0.61).

**Figure 5.**
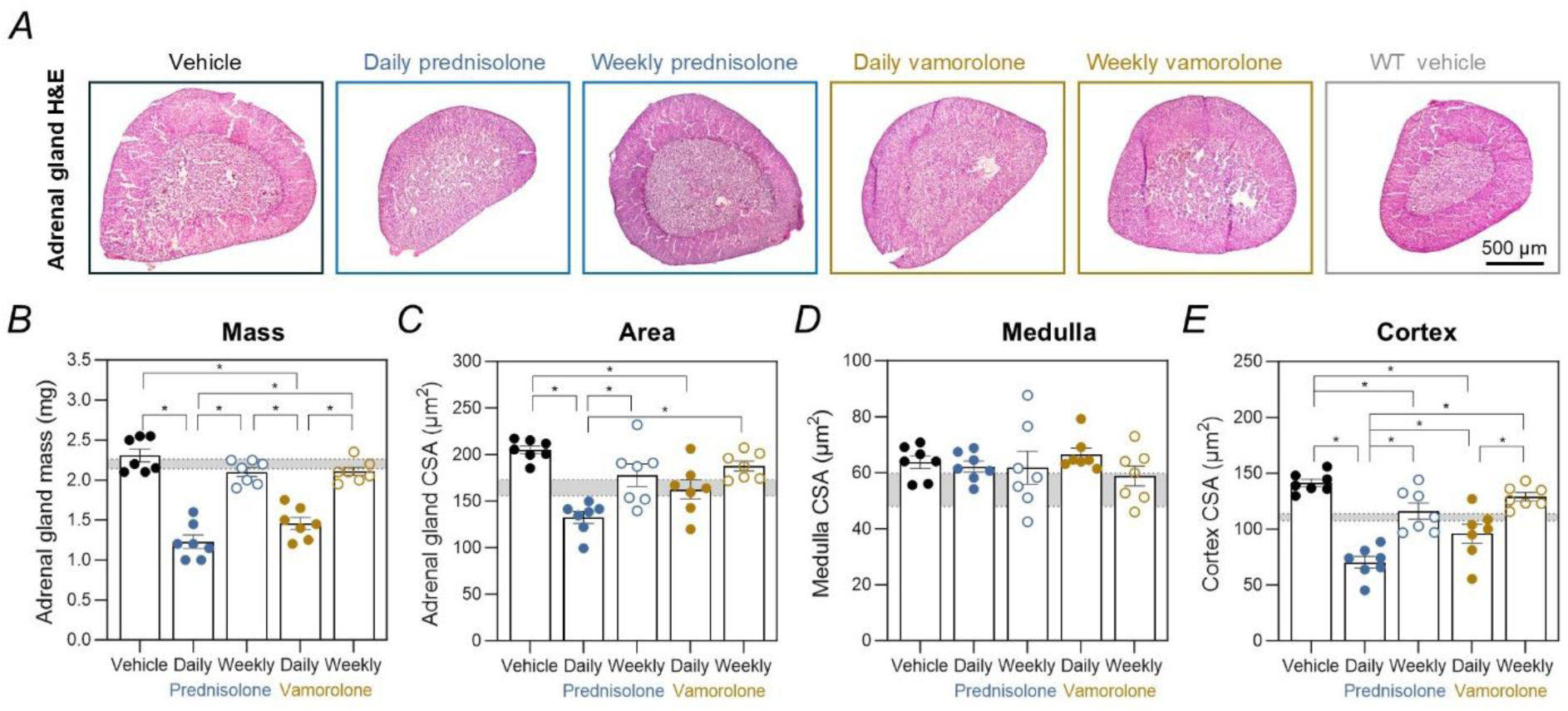
Daily glucocorticoid administration induces adrenal cortical atrophy, with a more pronounced effect following prednisolone compared with vamorolone. (A) Representative haematoxylin and eosin (H&E)-stained sections of adrenal glands (scale bar = 500 µm). (B) Adrenal gland mass. (C) Adrenal gland cross-sectional area (CSA). (D) Adrenal medulla CSA. (E) Adrenal cortex CSA. Group differences were assessed using univariate ANOVA; *P* < 0.05. WT responses are shown in grey shading as ± SEM. All other data are presented as mean ± SEM; *n* = 7 mice per group.

### 3.5 Cardiovascular physiology remains largely stable across glucocorticoid treatments

To assess treatment-dependent effects on cardiovascular physiology, we evaluated cardiac mass, haemodynamics, and peripheral perfusion. NMDS was then performed to integrate outcomes from mean arterial pressure, heart rate, tail blood flow, and heart mass into a single composite analysis. The NMDS showed that prednisolone, independent of dosing, did not alter overall heart physiology relative to *mdx* vehicle controls (P ≥ 0.053), however daily vamorolone did have an effect (**Figure 6A,B**; P = 0.037). Both daily and weekly prednisolone-treated mice were also comparable to WT controls (P ≥ 0.28), unlike vamorolone- or vehicle-treated *mdx* mice (**Figure 6A,B**; P ≥ 0.053). Daily vamorolone was associated with greater heart mass than prednisolone-treated mice, although no treatment differed from vehicle controls **(Figure 6C**; **Figure S5A**; P ≤ 0.016). Individually, treatment had no detectable effect on mean arterial pressure (P ≥ 0.86), heart rate (P ≥ 0.78), shock index (P ≥ 0.31), or tail blood flow (**Figure 6D-G**; P ≥ 0.22), demonstrating the strength of integrated multivariate analysis in revealing subtle cardiovascular effects that single metrics alone cannot detect.

**Figure 6.**
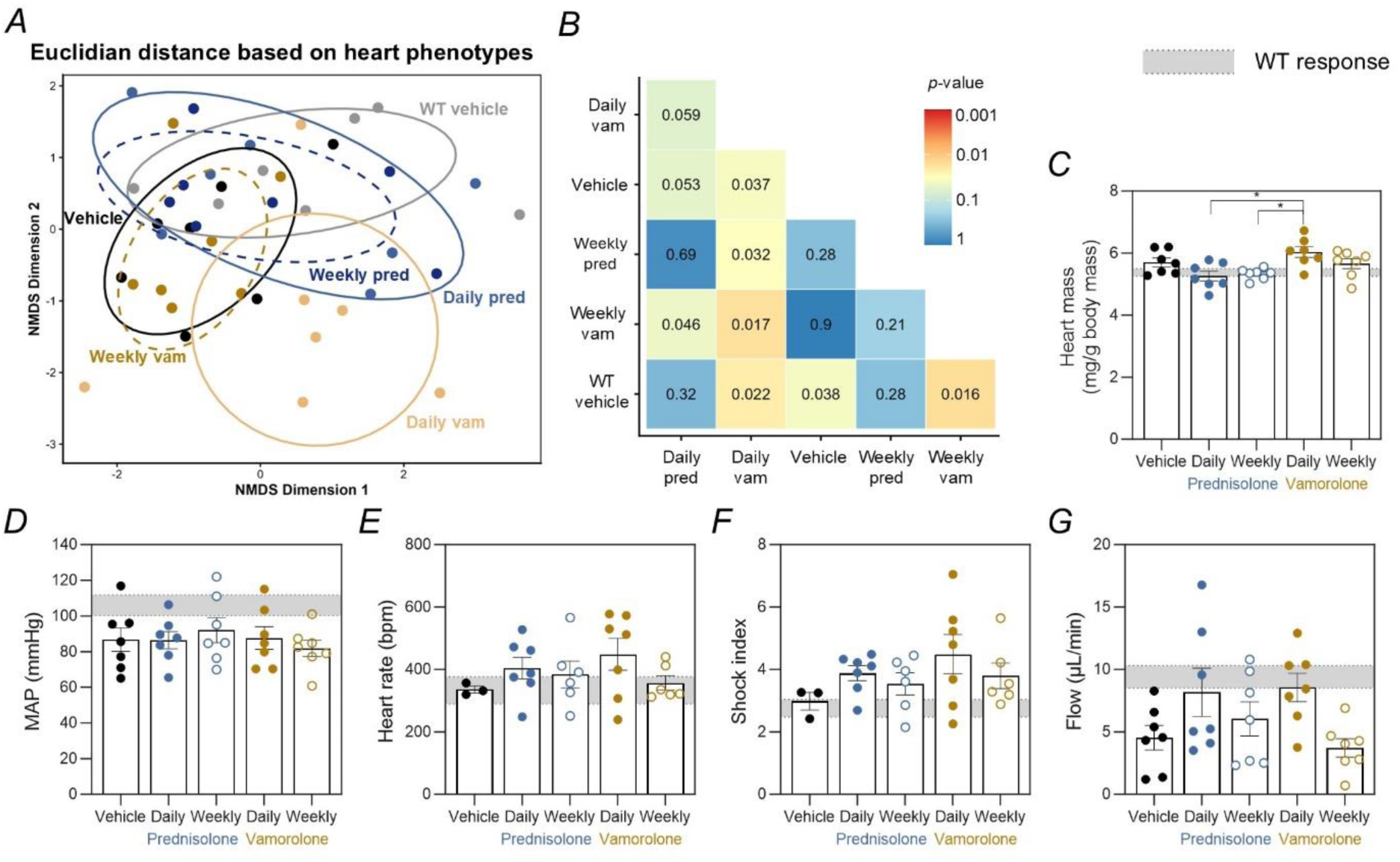
Striated muscle–associated cardiovascular physiology remains largely stable across glucocorticoid treatments, with limited modulation restricted to daily vamorolone administration. (A) Non-metric multidimensional scaling (NMDS) ordination based on Euclidean distance illustrating multivariate striated muscle physiological profiles across experimental groups, including prednisolone (blue) and vamorolone (yellow) treatments, and WT (black) and *mdx* (grey) vehicle controls. Each point represents an individual animal, and proximity reflects similarity in integrated physiological profiles. (B) Heat map of pairwise *p*-values for between-group multivariate comparisons across all experimental groups. (C) Heart mass expressed relative to body mass. (D) Mean arterial pressure (MAP), (E) heart rate, (F) shock index (maximum heart rate/low systolic blood pressure; index of hypovolemic shock) and (G) tail blood flow. Multivariate group differences were assessed using Euclidean distance–based PERMANOVA on z-score standardised data, with dispersion tested by betadisper. Significant global effects were followed by pairwise multivariate comparisons, and outcome-specific differences were assessed using univariate ANOVA with appropriate correction for multiple testing; *P* < 0.05. WT responses are shown in grey shading as ± SEM. All other data are presented as mean ± SEM; *n* = 7 mice per group.

### 3.6 Daily glucocorticoid administration disrupts systemic metabolic physiology and body composition

We next assessed systemic metabolic function to determine how GC treatment regimens broadly impact whole-body energy balance and organ physiology. Treating *mdx* mice during a juvenile growth window to model paediatric DMD, we observed that daily GCs impaired body mass growth (P ≤ 0.034) while paradoxically increasing food intake (P ≤ 0.042) relative to vehicle controls (**Figure 7A-C)**. Both daily prednisolone and daily vamorolone reduced fasting (P ≤ 0.019) and non-fasting (P ≤ 0.004) blood glucose levels relative to *mdx* vehicle controls, with no differences between these two daily treatments **Figure 7D,E**; P ≥ 0.286). Daily dosing of GCs decreased spleen mass (P ≤ 0.0001), although daily prednisolone caused a greater reduction than daily vamorolone (**Figure 7F**; **Figure S5B**; P = 0.0008). Weekly dosing of prednisolone (P = 0.012) but not vamorolone (P = 0.430) caused a reduction in spleen mass relative to vehicle controls (**Figure 7F)**. Daily GCs reduced long bone robustness (bone mass per unit length, mg/mm) relative to *mdx* vehicle controls (P ≤ 0.0006), while weekly dosing produced milder but still measurable decreases in the femur (**Figure 7G,H**; **Figure S5C-F**; P ≤ 0.002). Overall, these findings demonstrate that daily GC exposure imposes broad metabolic and systemic effects compromising growth, energy balance, immune organ size, and bone integrity; effects that are largely mitigated by weekly dosing.

**Figure 7.**
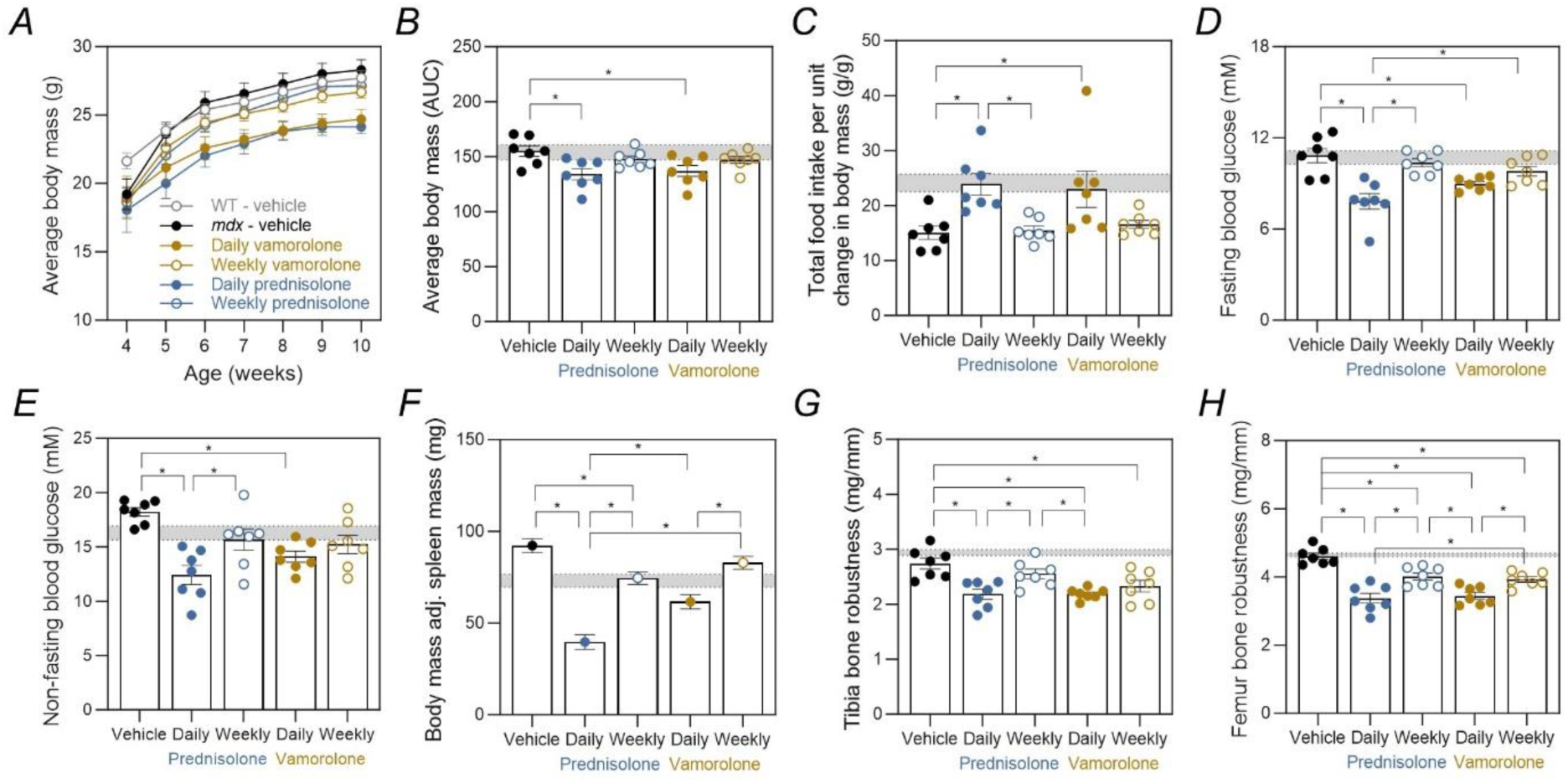
Daily glucocorticoids impair growth and body composition, causing hyperphagia, hypoglycaemia, and reduced bone robustness, with spleen mass decreased only by daily prednisolone. (A) Weekly body mass trajectories across the treatment period. (B) Body mass area under the curve (AUC). (C) Total food intake normalised to change in body mass over the treatment period. (D) Fasting blood glucose following 2 weeks of treatment. (E) Non-fasting blood glucose following 6 weeks of treatment. (F) Body-mass–adjusted spleen mass across treatment groups. Data are estimated marginal means (± SEM) derived from an ANCOVA model with body mass included as a covariate. (G) Tibial bone robustness (mass/length). (H) Femoral bone robustness (mass/length). Group differences were assessed using univariate ANOVA; *P* < 0.05. WT responses are shown in grey shading as ± SEM. All other data, except where otherwise specified, are presented as mean ± SEM; *n* = 7 mice per group.

## 4 Discussion

We investigated how GC treatment affects striated muscle, systemic metabolism, and CNS outcomes in juvenile *mdx* mice, comparing daily and weekly dosing of prednisolone and vamorolone over six weeks. Our results reveal that changes in striated muscle structure and function are mechanistically distinct and dosing regimen dependent, and that peripheral endocrine changes are largely uncoupled from CNS stress and anxiety responses. Weekly-dosed vamorolone produced the greatest gains in functional and *in vivo* strength, a notable finding, as previous studies of intermittent GC regimens (prednisolone and deflazacort) have not reported improvements in mass-adjusted strength parameters (Quattrocelli *et al*., 2017). In contrast, daily prednisolone had minimal impact on torque generation despite reducing circulating CK and driving muscle microstructural remodelling. System-level NMDS analyses captured coordinated effects, highlighting compound- and regimen-specific trade-offs between muscle efficacy and systemic tolerability, including adrenal, metabolic, and bone phenotypes, and revealing patterns not apparent from individual measures alone.

The relationship between striated muscle microstructure and functional performance is complex (Charles *et al*., 2022), and here we show that GC treatments appear to dissociate structural remodelling from strength outcomes in *mdx* mice. Daily prednisolone reduced myofibre CSA and increased fibre density (number of fibres per unit area). Myofibres from vamorolone-treated mice maintained the same fibre density despite smaller CSA. This less disruptive remodelling pattern may reflect vamorolone’s reduced catabolic effects compared to traditional GCs, consistent with its dissociative steroid mechanism (Heier *et al*., 2019; Crastin *et al*., 2025). Weekly dosing mitigated these structural effects across both compounds, consistent with previous reports that daily GC exposure triggers atrophy pathways and impairs voluntary muscle performance, whereas intermittent dosing avoids atrophy and preserves or improves muscle function (Quattrocelli *et al*., 2017). However, while prednisolone induced more pronounced myofibre remodelling relative to vamorolone, these structural changes did not correlate with functional strength outcomes in either compound, highlighting how GC-induced muscle fibre remodelling can occur independently of muscle performance in *mdx* mice.

Limiting ongoing muscle damage is a critical goal in DMD, as repeated myofibre injury drives progressive weakness and disease progression (Duan *et al*., 2021). Daily prednisolone was the only regimen to reduce circulating CK, indicating meaningful protection against ongoing muscle damage. However, weekly dosing alone is not enough to mitigate damage given that weekly prednisolone was insufficient to reduce CK. Notably, these findings contrast with reports that vamorolone acts as a more potent membrane stabiliser than prednisolone and thus should theoretically produce even lower CK elevations (Heier *et al*., 2013). Although CK reduction did not translate to short-term strength gains, this represents a disease-modifying effect that may confer longer-term functional benefit beyond the scope of this study.

While the primary physiological benefits of GC treatment in improving muscle function in DMD has been investigated (Baltgalvis *et al*., 2009; Guerron *et al*., 2010; Sali *et al*., 2012; Ricotti *et al*., 2013; Bello *et al*., 2015; Morrison-Nozik *et al*., 2015; Quattrocelli *et al*., 2017, 2019; McDonald *et al*., 2018), much less is known about their neurocognitive and endocrine consequences. Using an integrative analysis, we confirm that *mdx* mice exhibit distinct stress and anxiety physiology relative to WT mice, yet GCs neither exacerbate nor alleviate these CNS dysfunctions. Limited GC penetration (Liu *et al*., 2024) or intrinsic GABAergic deficits (Vaillend & Chaussenot, 2017) in *mdx* mice may explain the relative insensitivity of these circuits. Supporting the latter, stress- and anxiety-related behaviours remained unchanged despite clear adrenal cortical atrophy with daily dosing, demonstrating that central stress phenotypes are not simply driven by peripheral HPA-axis output (Lindsay & Russell, 2023). Notably, adrenal atrophy was most pronounced with prednisolone, whereas vamorolone, particularly when dosed weekly, largely spared adrenal structure, helping to reconcile heterogeneity in reported adrenal suppression across preclinical and clinical studies (Heier *et al*., 2013; Conklin *et al*., 2018; Guglieri *et al*., 2022; Liu *et al*., 2024; Dang *et al*., 2024). Together, these findings indicate that differential peripheral endocrine suppression does not translate into exacerbated CNS-related phenotypes.

Cardiovascular physiology remained stable across treatments, reflecting minimal overt cardiac pathology in juvenile *mdx* mice (Adamo *et al*., 2010; Verhaart *et al*., 2012; Wasala *et al*., 2013). Daily vamorolone uniquely increased normalised heart mass relative to prednisolone-treated mice and drove subtle NMDS shifts away from WT controls, despite no individual haemodynamic changes, raising the possibility of early pathological remodelling despite previously predicted cardioprotection via MR antagonism (Heier *et al*., 2019). However, while juvenile *mdx* models best replicate early DMD skeletal muscle pathology, cardiovascular stability limits their utility for long-term cardiac risk assessment, as echocardiographic defects emerge only after 10 months (Yucel *et al*., 2018).

Daily GC treatment imposed substantial systemic adverse effects in *mdx* mice, largely independent of muscle-specific efficacy. Mice receiving daily GCs exhibited suppressed growth, reduced metabolic throughput, decreased spleen mass (prednisolone only), and compromised long bone robustness. Hyperphagia accompanied by low circulating glucose levels indicated inefficient substrate utilisation (Quattrocelli *et al*., 2019), consistent with a GC-induced hypometabolic state characterised by suppressed growth (Quattrocelli *et al*., 2017), increased catabolism, reduced glucose production, and impaired fasting-induced glucose mobilisation. In contrast, weekly dosing, particularly of vamorolone, attenuated these systemic effects, highlighting dosing frequency as a key determinant of therapeutic index. These findings demonstrate that the systemic adverse effect profile is dose-dependent and that careful optimisation of GC regimens can mitigate adverse outcomes while retaining potential therapeutic benefit.

This study was limited to juvenile *mdx* mice (4–10 weeks), which capture early muscle degeneration but not long-term DMD progression, late cardiac pathology, or chronic GC effects. The six-week duration and single doses per regimen also restrict insights into sustained efficacy and dose optimisation, while modest CNS sample sizes may have obscured subtle neurobehavioral changes. Future studies should not only address these limitations but also elucidate the mechanistic basis for the observed structure-function dissociation and how optimised regimens reshape muscle regeneration at the cellular level.

Taken together, our findings demonstrate that therapeutic efficacy in DMD cannot be defined solely by muscle-specific outcomes. Daily GCs provided meaningful structural and damage-limiting effects in muscle but imposed substantial systemic costs, including growth suppression, metabolic hypofunction, adrenal atrophy, and reduced long bone robustness. Intermittent dosing or dissociative compounds, demonstrated by weekly vamorolone, improved muscle functional performance while mitigating many of these adverse effects, highlighting the critical role of dosing frequency - a first systematic investigation for vamorolone and for higher dose prednisolone. These results underscore the value of multi-system integrative analyses to define the therapeutic index, providing a framework to optimise GC therapy. Given that GCs remain the only widely available therapy for all boys with DMD, these findings highlight the value in continued optimisation of GC dosing, both for classical and dissociative steroids, to maximise muscle benefits while minimising systemic side effects, supporting safer and more effective long-term treatment for boys with DMD.

## Supplementary Figures

**Figure S1.**
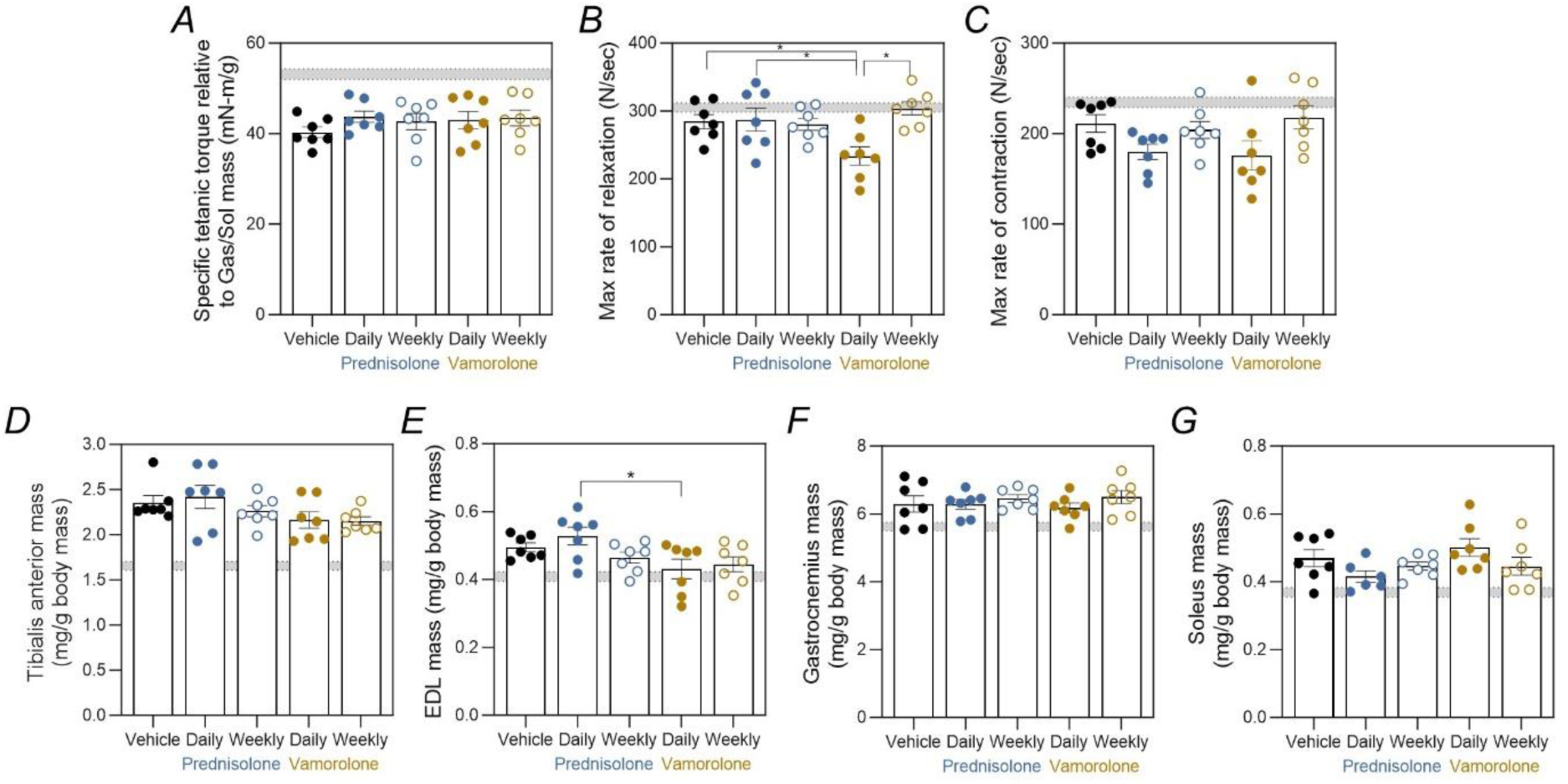
Evaluation of posterior leg muscle strength, contractile properties, and relative muscle mass. (A) *In vivo* muscle strength expressed as specific tetanic torque normalised to gastrocnemius (gas) and soleus (sol) muscle mass. (B) Contraction kinetics of posterior leg muscles expressed as maximal rate of torque development. (C) Relaxation kinetics of posterior leg muscles expressed as maximal rate of torque relaxation. (D) Tibialis anterior mass expressed relative to body mass. (E) EDL muscle mass expressed relative to body mass. (F) Gastrocnemius mass expressed relative to body mass. (G) Soleus mass expressed relative to body mass. Group differences were assessed using univariate ANOVA; *P* < 0.05. WT responses are shown in grey shading as ± SEM. All other data are presented as mean ± SEM; *n* = 7 mice per group.

**Figure S2.**
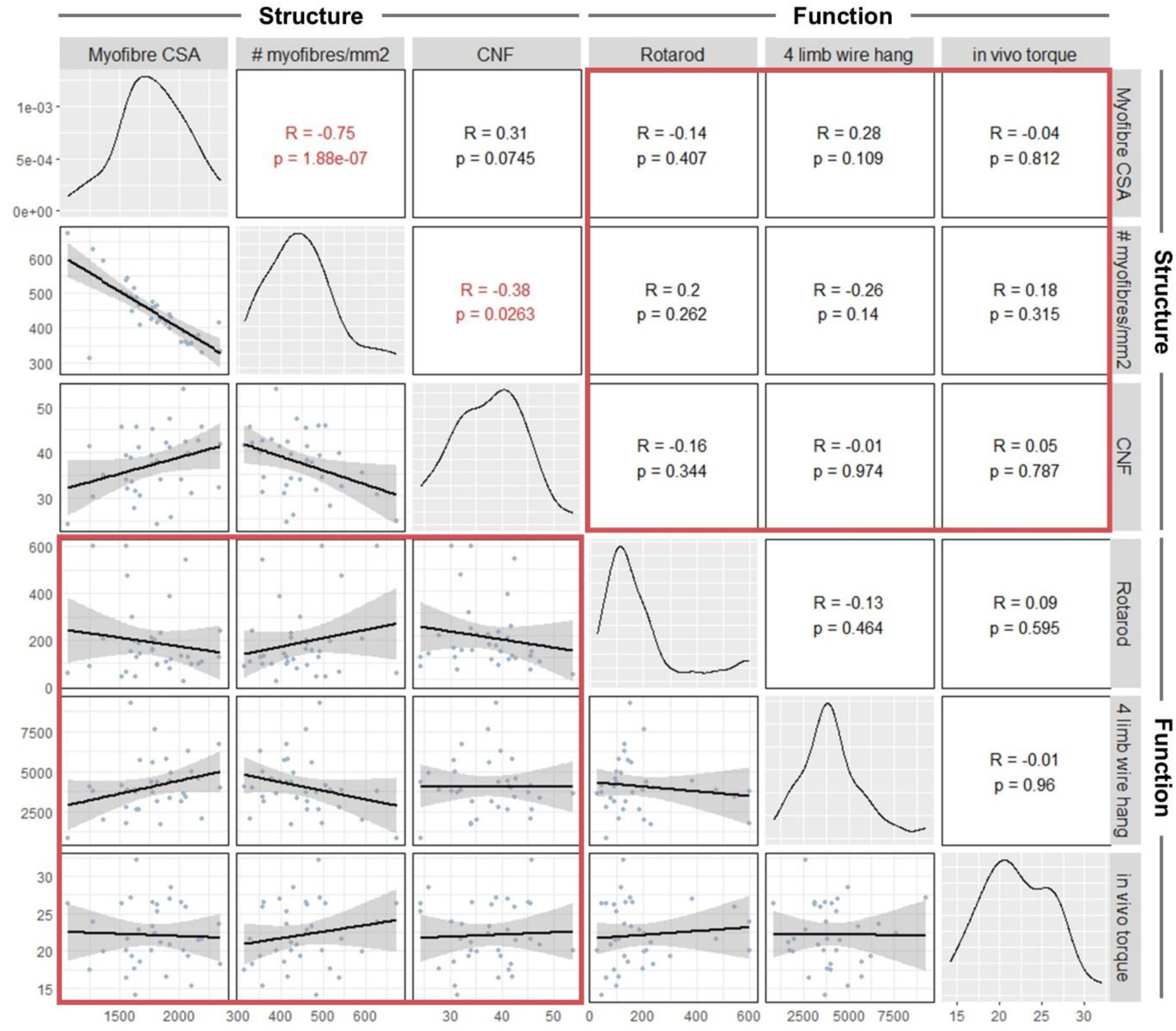
Relationship between muscle structural and functional parameters in *mdx* muscle. Pairwise scatterplots showing associations between architectural muscle fibre characteristics (myofibre cross-sectional area, fibre number density, and proportion of centronucleated fibres) and functional outcomes (rotarod performance, four-limb wire hang, and *in vivo* torque) in *mdx* mice. Lower panels show individual data points with linear regression fits and 95% confidence intervals; upper panels show Pearson correlation coefficients (R) and corresponding *p*-values.

**Figure S3.**
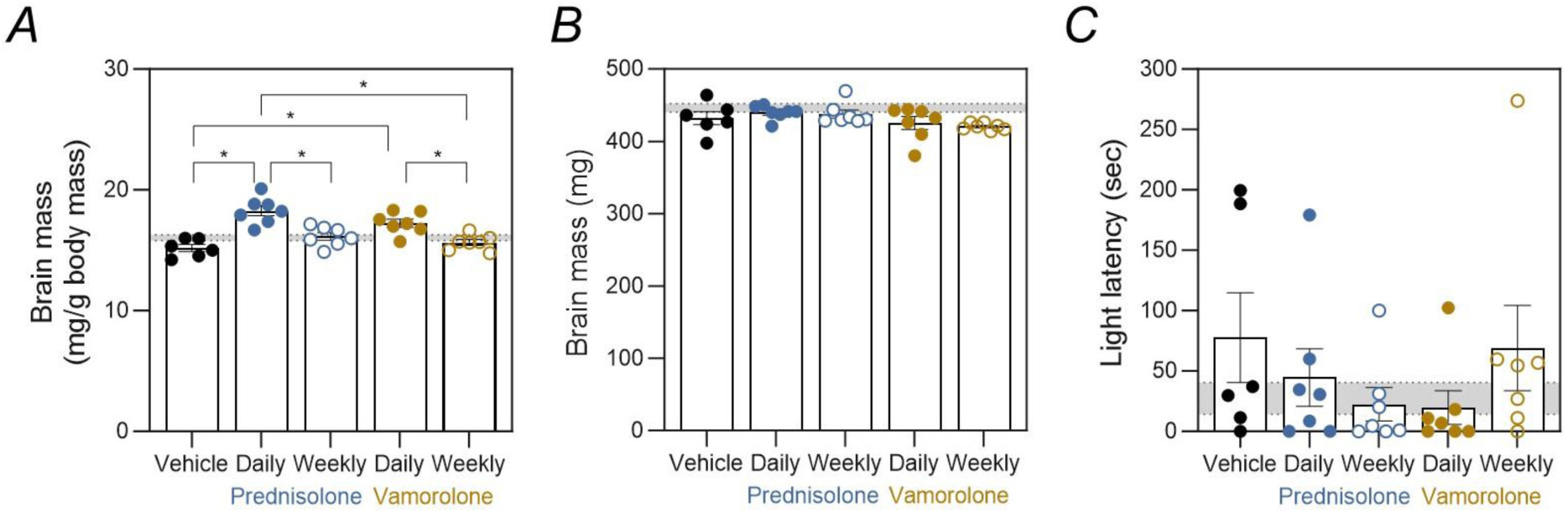
Assessment of brain size and anxiety-related behaviour. (A) Brain mass expressed relative to body mass. (B) Absolute brain mass. *Light–dark choice anxiety test:* (C) Latency to enter lit compartment. Group differences were assessed using univariate ANOVA or Kruskall-Wallis H Test; *P* < 0.05. WT responses are shown in grey shading as ± SEM. All other data are presented as mean ± SEM; *n* = 7 mice per group.

**Figure S4.**
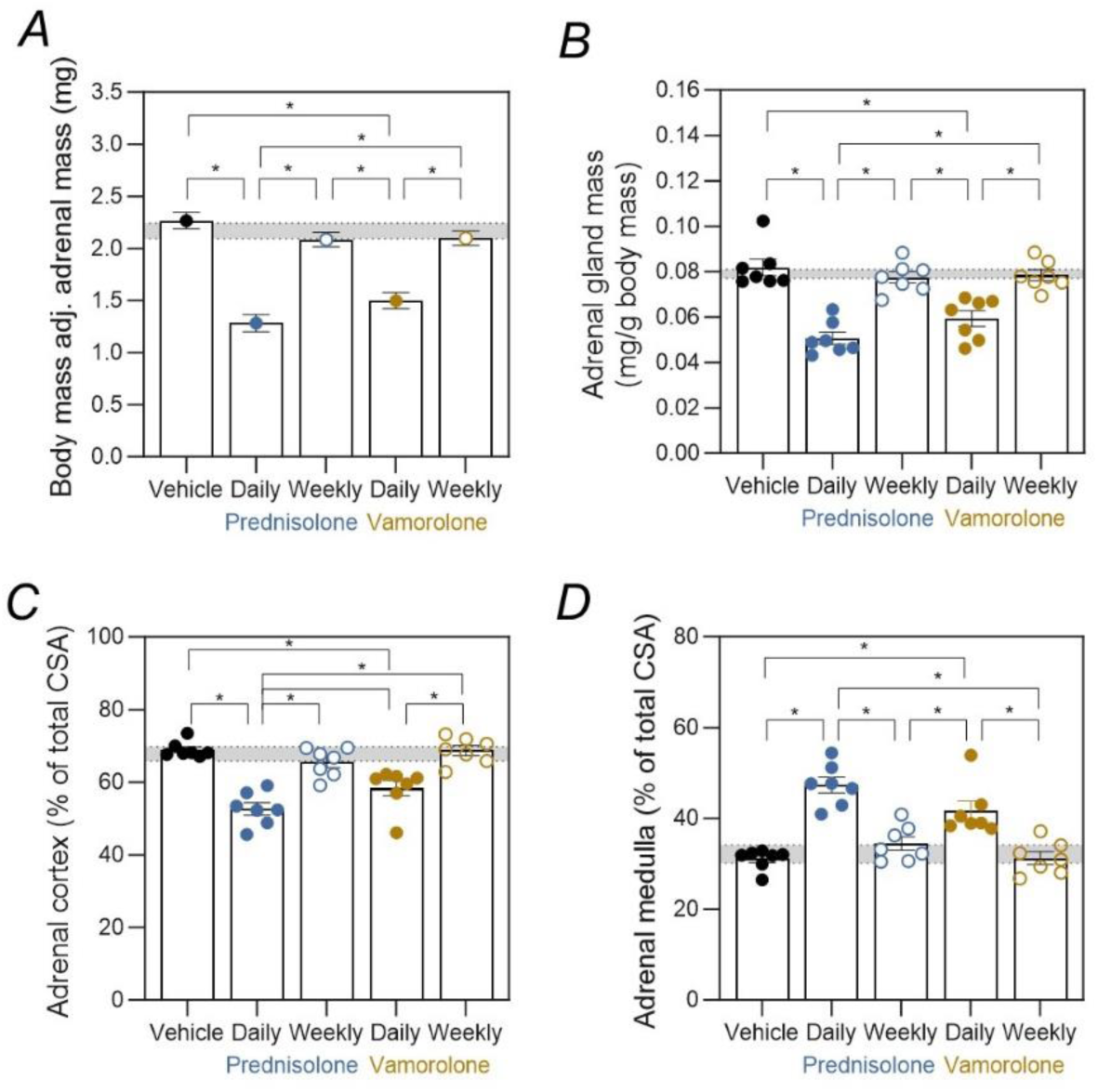
Overview of adrenal gland size and regional composition. (A) Body-weight–adjusted adrenal gland mass across treatment groups. Data are estimated marginal means (± SEM) derived from an ANCOVA model with body weight included as a covariate. (B) Adrenal gland mass expressed relative to body mass. (C) Adrenal gland cortex area expressed as a percentage of adrenal gland cross sectional area (CSA). (D) Adrenal gland medulla area expressed as a percentage of adrenal gland CSA. Note that the apparent increase in medullary percentage reflects cortical atrophy rather than genuine medullary enlargement, as absolute medulla CSA was unchanged across groups. Group differences were assessed using univariate ANOVA; *P* < 0.05. WT responses are shown in grey shading as ± SEM. All other data, except where otherwise specified, are presented as mean ± SEM; *n* = 7 mice per group.

**Figure S5.**
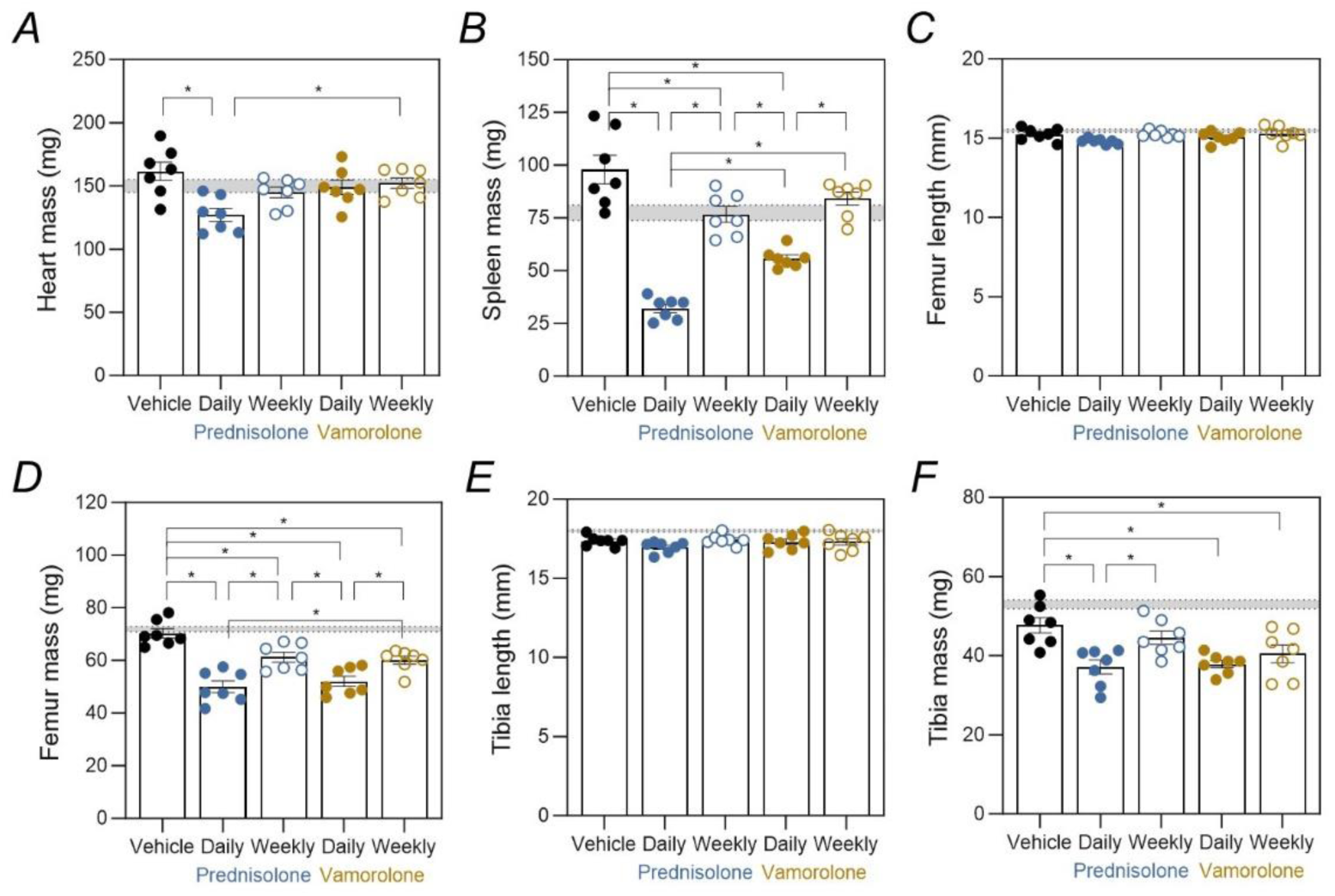
Overview of organ and bone size. (A) Heart mass. (B) Spleen mass. (C) Femoral length. (D) Femoral mass. (E) Tibial length. (F) Tibial mass. Group differences were assessed using univariate ANOVA; *P* < 0.05. WT responses are shown in grey shading as ± SEM. All other data are presented as mean ± SEM; *n* = 7 mice per group.

## References

Adamo CM, Dai DF, Percival JM, Minami E, Willis MS, Patrucco E, Froehner SC & Beavo JA (2010). Sildenafil reverses cardiac dysfunction in the mdx mouse model of Duchenne muscular dystrophy. Proc Natl Acad Sci U S A 107, 19079–19083.

Angelini C & Peterle E (2012). Old and new therapeutic developments in steroid treatment in Duchenne muscular dystrophy. Acta Myologica 31, 9–15.

Baltgalvis KA, Call JA, Nikas JB & Lowe DA (2009). Effects of prednisolone on skeletal muscle contractility in mdx mice. Muscle Nerve 40, 443–454.

Bello L, Gordish-Dressman H, Morgenroth LP, Henricson EK, Duong T, Hoffman EP, Cnaan A & Mcdonald CM (2015). Prednisone/prednisolone and deflazacort regimens in the CINRG Duchenne Natural History Study. Neurology 85, 1048–1055.

Caudal D, François V, Lafoux A, Ledevin M, Anegon I, Le Guiner C, Larcher T & Huchet C (2020). Characterization of brain dystrophins absence and impact in dystrophin-deficient Dmdmdx rat model. PLoS One 15, e0230083.

Charles J, Kissane R, Hoehfurtner T & Bates KT (2022). From fibre to function: are we accurately representing muscle architecture and performance? Biological Reviews 97, 1640–1676.

Conklin LS et al. (2018). Phase IIa trial in Duchenne muscular dystrophy shows vamorolone is a first-in-class dissociative steroidal anti-inflammatory drug. Pharmacol Res 136, 140–150.

Connolly AM, Schierbecker J, Renna R & Florence J (2002). High dose weekly oral prednisone improves strength in boys with Duchenne muscular dystrophy. Neuromuscular disorders 12, 917–925.

Crastin A, Shanker A, Sagmeister MS, Taylor A, Lavery GG, Raza K & Hardy RS (2025). Vamorolone: a novel metabolism resistant steroid that suppresses joint destruction in chronic polyarthritis with reduced systemic side effects. Rheumatology 64, 4371–4381.

Dang UJ et al. (2024). Efficacy and safety of Vamorolone over 48 weeks in boys with Duchenne muscular dystrophy. Neurology 102, e208112.

Darmahkasih AJ, Rybalsky I, Tian C, Shellenbarger KC, Horn PS, Lambert JT & Wong BL (2020). Neurodevelopmental, behavioral, and emotional symptoms common in Duchenne muscular dystrophy. Muscle Nerve 61, 466–474.

Devananthan P, Craven R, Joe K, Major GS, Chen J, Kabaliuk N & Lindsay A (2026). Dystrophin deficiency stiffens skeletal muscle and impairs elasticity: an in vivo rheological examination. J Appl Physiol 140, 39–56.

Duan D, Goemans N, Takeda S, Mercuri E & Aartsma-Rus A (2021). Duchenne muscular dystrophy. Nat Rev Dis Primers; DOI: 10.1038/s41572-021-00248-3.

Ferreira-Duarte M, Lopes IM, Morato M & Duarte-Araújo M (2024). Rats prefer condensed milk to strawberry jam – a new possibility for voluntary oral drug administration. Lab Anim 58, 160–163.

Gharibi S, Major GS, Shad A, Semple BD, McGregor NE, Blank M, Abbott G, Sims NA, Shaw CS, Russell AP & Lindsay A (2025). Stress-induced fetal programming contributes to the manifestation of Duchenne muscular dystrophy in mdx mice. iScience; DOI: 10.1016/j.isci.2025.112123.

Gharibi S, Vaillend C & Lindsay A (2024). The unconditioned fear response in vertebrates deficient in dystrophin. Prog Neurobiol 235, 102590.

Goldstein JA & McNally EM (2010). Mechanisms of muscle weakness in muscular dystrophy. In Journal of General Physiology, pp. 29–34.

Guerron AD, Rawat R, Sali A, Spurney CF, Pistilli E, Cha HJ, Pandey GS, Gernapudi R, Francia D, Farajian V, Escolar DM, Bossi L, Becker M, Zerr P, de la Porte S, Gordish-Dressman H, Partridge T, Hoffman EP & Nagaraju K (2010). Functional and molecular effects of arginine butyrate and prednisone on muscle and heart in the mdx mouse model of Duchenne muscular dystrophy. PLoS One; DOI: 10.1371/journal.pone.0011220.

Guglieri M et al. (2022). Efficacy and safety of Vamorolone vs placebo and Prednisone among boys with Duchenne muscular dystrophy: A randomized clinical trial. JAMA Neurol 79, 1005–1014.

Handberg C, Werlauff U & Højberg AL (2022). Perspectives on everyday life challenges of Danish young people with Duchenne muscular dystrophy (DMD) on corticosteroids. Glob Qual Nurs Res; DOI: 10.1177/23333936221094858.

Heier CR et al. (2013). VBP15, a novel anti-inflammatory and membrane-stabilizer, improves muscular dystrophy without side effects. EMBO Mol Med 5, 1569–1585.

Heier CR, Yu Q, Fiorillo AA, Tully CB, Tucker A, Mazala DA, Uaesoontrachoon K, Srinivassane S, Damsker JM, Hoffman EP, Nagaraju K & Spurney CF (2019). Vamorolone targets dual nuclear receptors to treat inflammation and dystrophic cardiomyopathy. Life Sci Alliance; DOI: 10.26508/LSA.201800186.

Hoffman EP, Brown RH & Kunkel LM (1987). Dystrophin: the protein product of the Duchenne muscular dystrophy locus. Cell 51, 919–928.

Hughes DC, Marcotte GR, Marshall AG, West DWD, Baehr LM, Wallace MA, Saleh PM, Bodine SC & Baar K (2017). Age-related differences in dystrophin: Impact on force transfer proteins, membrane integrity, and neuromuscular junction stability. Journals of Gerontology - Series A Biological Sciences and Medical Sciences 72, 640–648.

Kourakis S, Timpani CA, Bagaric RM, Qi B, Ali BA, Boyer R, Spiesberger G, Kandhari N, Peterson AL, Debrincat D, Yates TJ, Yan X, Kuang J, de Haan JB, Stupka N, Nijagal B, Deveson-Lucas D, Fischer D & Rybalka E (2024). Moderate-term dimethyl fumarate treatment reduces pathology of dystrophic skeletal and cardiac muscle in a mouse model. ; DOI: 10.1101/2024.07.13.601627. Available at: http://biorxiv.org/lookup/doi/10.1101/2024.07.13.601627.

Lindsay A, Holm J, Razzoli M, Bartolomucci A, Ervasti JM & Lowe DA (2021a). Some dystrophy phenotypes of dystrophin-deficient mdx mice are exacerbated by mild, repetitive daily stress. FASEB Journal 35, e21489.

Lindsay A & Russell AP (2023). The unconditioned fear response in dystrophin-deficient mice is associated with adrenal and vascular function. Sci Rep 13, 55513.

Lindsay A, Trewin AJ, Sadler KJ, Laird C, Della Gatta PA & Russell AP (2021b). Sensitivity to behavioral stress impacts disease pathogenesis in dystrophin-deficient mice. FASEB Journal 35, e22034.

Liu G et al. (2024). Comparison of pharmaceutical properties and biological activities of prednisolone, deflazacort, and vamorolone in DMD disease models. Hum Mol Genet 33, 211–223.

Maresh K, Papageorgiou A, Ridout D, Harrison NA, Mandy W, Skuse D & Muntoni F (2023). Startle responses in Duchenne muscular dystrophy: a novel biomarker of brain dystrophin deficiency. Brain 146, 252–265.

McCormack NM, Nguyen NY, Tully CB, Oliver T, Fiorillo AA & Heier CR (2023). Vamorolone improves Becker muscular dystrophy and increases dystrophin protein in bmx model mice. iScience; DOI: 10.1016/j.isci.2023.107161.

McDonald CM et al. (2018). Long-term effects of glucocorticoids on function, quality of life, and survival in patients with Duchenne muscular dystrophy. The Lancet 391, 451–461.

Mochizuki H, Miyatake S, Suzuki M, Shigeyama T, Yatabe K, Ogata K, Tamura T & Kawai M (2008). Mental retardation and lifetime events of Duchenne muscular dystrophy in Japan. Internal Medicine 47, 1207–1210.

Morrison-Nozik A, Anand P, Zhu H, Duan Q, Sabeh M, Prosdocimo DA, Lemieux ME, Nordsborg N, Russell AP, MacRae CA, Gerber AN, Jain MK & Haldar SM (2015). Glucocorticoids enhance muscle endurance and ameliorate DMD through a defined metabolic program. Proc Natl Acad Sci U S A 112, E6780–E6789.

Quattrocelli M, Barefield DY, Warner JL, Vo AH, Hadhazy M, Earley JU, Demonbreun AR & McNally EM (2017). Intermittent glucocorticoid steroid dosing enhances muscle repair without eliciting muscle atrophy. Journal of Clinical Investigation 127, 2418–2432.

Quattrocelli M, Zelikovich AS, Jiang Z, Peek CB, Demonbreun AR, Kuntz NL, Barish GD, Haldar SM, Bass J & McNally EM (2019). Pulsed glucocorticoids enhance dystrophic muscle performance through epigenetic-metabolic reprogramming. JCI Insight 4, e132402.

Razzoli M, Lindsay A, Law ML, Chamberlain CM, Southern WM, Berg M, Osborn J, Engeland WC, Metzger JM, Ervasti JM & Bartolomucci A (2020). Social stress is lethal in the mdx model of Duchenne muscular dystrophy. EBioMedicine 55, 02700.

Ricotti V et al. (2013). Long-term benefits and adverse effects of intermittent versus daily glucocorticoids in boys with Duchenne muscular dystrophy. J Neurol Neurosurg Psychiatry 84, 698–705.

Sali A, Guerron AD, Gordish-Dressman H, Spurney CF, Iantorno M, Hoffman EP & Nagaraju K (2012). Glucocorticoid-treated mice are an inappropriate positive control for long-term preclinical studies in the mdx mouse. PLoS One 7, e34204.

Saoudi A, Zarrouki F, Sebrié C, Izabelle C, Goyenvalle A & Vaillend C (2021). Emotional behavior and brain anatomy of the mdx52 mouse model of Duchenne muscular dystrophy. DMM Disease Models and Mechanisms; DOI: 10.1242/dmm.049028.

Scarborough J, Mueller F, Arban R, Dorner-Ciossek C, Weber-Stadlbauer U, Rosenbrock H, Meyer U & Richetto J (2020). Preclinical validation of the micropipette-guided drug administration (MDA) method in the maternal immune activation model of neurodevelopmental disorders. Brain Behav Immun 88, 461–470.

Sekiguchi M, Zushida K, Yoshida M, Maekawa M, Kamichi S, Yoshida M, Sahara Y, Yuasa S, Takeda S & Wada K (2009). A deficit of brain dystrophin impairs specific amygdala GABAergic transmission and enhances defensive behaviour in mice. Brain 132, 124–135.

du Sert NP et al. (2020). The arrive guidelines 2.0: Updated guidelines for reporting animal research. PLoS Biol 18, e3000410.

Smith LR & Barton ER (2014). SMASH - semi-automatic muscle analysis using segmentation of histology: A MATLAB application. Skelet Muscle 4, 21.

Timpani CA et al. (2023). Dimethyl fumarate modulates the dystrophic disease program following short-term treatment. ; DOI: 10.1172/jci.

Ullrich ND, Fanchaouy M, Gusev K, Shirokova N & Niggli E (2009). Hypersensitivity of excitation-contraction coupling in dystrophic cardiomyocytes. Am J Physiol Heart Circ Physiol; DOI: 10.1152/ajpheart.00602.2009.

Vaillend C, Aoki Y, Mercuri E, Hendriksen J, Tetorou K, Goyenvalle A & Muntoni F (2025). Duchenne muscular dystrophy: recent insights in brain related comorbidities. Nat Commun 16, 1298.

Vaillend C & Chaussenot R (2017). Relationships linking emotional, motor, cognitive and GABAergic dysfunctions in dystrophin-deficient mdx mice. Hum Mol Genet 26, 1041–1055.

Verhaart IEC, van Duijn RJM, den Adel B, Roest AAW, Verschuuren JJGM, Aartsma-Rus A & van der Weerd L (2012). Assessment of cardiac function in three mouse dystrophinopathies by magnetic resonance imaging. Neuromuscular Disorders 22, 418–426.

Wasala NB, Bostick B, Yue Y & Duan D (2013). Exclusive skeletal muscle correction does not modulate dystrophic heart disease in the aged mdx model of Duchenne cardiomyopathy. Hum Mol Genet 22, 2634–2641.

Yucel N, Chang AC, Day JW, Rosenthal N & Blau HM (2018). Humanizing the mdx mouse model of DMD: the long and the short of it. NPJ Regen Med; DOI: 10.1038/s41536-018-0045-4.

Ziemba M, Barkhouse M, Uaesoontrachoon K, Giri M, Hathout Y, Dang UJ, Gordish-Dressman H, Nagaraju K & Hoffman EP (2021). Biomarker-focused multi-drug combination therapy and repurposing trial in mdx mice. PLoS One; DOI: 10.1371/journal.pone.0246507.

